# Comparative assessment of fluorescent proteins for *in vivo* imaging in an animal model system

**DOI:** 10.1101/040279

**Authors:** Jennifer K. Heppert, Daniel J. Dickinson, Ariel M. Pani, Christopher D. Higgins, Annette Steward, Julie Ahringer, Jeffrey R. Kuhn, Bob Goldstein

**Author notes:** Correspondence: Bob Goldstein.

## Abstract

Fluorescent protein tags are fundamental tools used to visualize gene products and analyze their dynamics *in vivo*. Recent advances in genome editing have enabled precise insertion of fluorescent protein tags into the genomes of diverse organisms. These advances expand the potential of *in vivo* imaging experiments, and they facilitate experimentation with new, bright, photostable fluorescent proteins. Most quantitative comparisons of the brightness and photostability of different fluorescent proteins have been made *in vitro*, removed from biological variables that govern their performance in cells or organisms. To address the gap we quantitatively assessed fluorescent protein properties *in vivo* in an animal model system. We generated transgenic *C. elegans* strains expressing green, yellow, or red fluorescent proteins in embryos, and we imaged embryos expressing different fluorescent proteins under the same conditions for direct comparison. We found that mNeonGreen was not bright *in vivo* as predicted based on *in vitro* data, but that mNeonGreen is a better tag than GFP for specific kinds of experiments, and we report on optimal red fluorescent proteins. These results identify ideal fluorescent proteins for imaging *in vivo* in *C. elegans* embryos, and they suggest good candidate fluorescent proteins to test in other animal model systems.

**Abbreviations**
FP: fluorescent protein
GFP: green fluorescent protein
mNG: monomeric neon green
mYPet: monomeric yellow fluorescent protein for energy transfer
CRISPR: clustered, regularly interspersed, short palindromic repeats

## Introduction

For more than two decades, cell and developmental biologists have used genetically-encoded fluorescent protein fusion tags to visualize proteins in living cells and organisms. Efforts to engineer and discover superior fluorescent proteins have resulted in variants with diverse emission wavelengths and photophysical properties (Tsien, 1998; Matz *et al.,* 1999; Shaner *et al.,* 2004; 2007; shcherbo *et al.,* 2009; Shaner *et al.,* 2013; Shaner, 2014). The color, brightness, and photostability of a fluorescent protein are critical parameters to consider for experiments in which proteins will be imaged *in vivo* (Shaner *et al.,* 2005; Davidson and Campbell, 2009; Shaner, 2014). However, most brightness and photostability measurements are made with purified fluorescent proteins *in vitro* (Shaner *et al.,* 2005). While this approach provides information about the intrinsic optical properties of each fluorescent protein, it does not replicate many of the conditions of an *in vivo,* biological system.

Historically, many methods used to express fluorescently tagged proteins resulted in non-physiological levels of proteins of interest, limiting the interpretation of some experiments (Huang *et al.,* 2000; Krestel *et al.,* 2004; Doyon *et al.,* 2011). However, genome engineering techniques based on the CRISPR/Cas9 system have recently made it possible to more precisely edit the genomes of diverse cell types and organisms (Doudna and Charpentier, 2014; Gilles and Averof, 2014; Harrison *et al.,* 2014; Hsu *et al.,* 2014; Peng *et al.,* 2014) and to routinely insert fluorescent protein tags into endogenous genomic loci in some organisms, as has long been standard in yeast (Dickinson *et al.,* 2013; Auer *et al*., 2014; Bassett *et al*., 2014; Ma *et al*., 2014; Paix *et al*., 2014; Xue *et al.,* 2014; Aida *et al.,* 2015; Dickinson *et al.,* 2015; Perry and Henry, 2015; Ratz *et al.,* 2015). With this technological advance comes an increase in need for information about the best fluorescent proteins to use for *in vivo* imaging studies. Fortunately, advances in genome editing techniques have also created an opportunity to close this gap in knowledge by facilitating the comparison of fluorescent proteins *in vivo*.

Our goal in this study was to make a systematic comparison of fluorescent proteins that would answer the question: What fluorescent protein should one use *in vivo* for a given experiment? A previous systematic analysis of fluorescent proteins performed in *Saccharomyces cerevisiae* revealed clear information about which tags to use *in vivo* in yeast (Lee *et al.,* 2013). Since that study, new fluorescent proteins have been characterized, including some reported to be brighter than GFP (Shaner *et al.,* 2013). Here we report direct comparisons of monomeric green (GFP, mNeonGreen - mNG), yellow (mYPet, mNG), and red (TagRFP-T, mRuby2, mCherry, mKate2) fluorescent proteins *in vivo,* in a multicellular animal model organism. We used CRISPR/Cas9-triggered homologous recombination in *C. elegans* to express the same transgene tagged with optimized versions of various fluorescent proteins from the same genomic locus. This allowed us to quantitatively compare the brightness and photostability of these fluorescent proteins in embryos imaged under typical experimental conditions.

Our findings provide quantitative data that are useful for choosing which fluorescent proteins to use for *in vivo* experiments in *C. elegans*. The results suggest a set of candidate fluorescent proteins for testing in other model systems, and more generally, they demonstrate the value of testing fluorescent protein performance *in vivo*. We also contribute novel tools for the field including constructs containing optimized fluorescent proteins and an Excel based tool to assist investigators in choosing the best fluorescent proteins to use with their imaging resources.

## Results and Discussion

### Generating single-copy transgene knock-ins

To directly compare fluorescent proteins *in vivo,* we used CRIPSR/Cas-9 to generate single-copy transgene knock-in strains expressing distinct fluorescent proteins. Constructs used to create these strains were identical except for the fluorescent protein sequences encoded in each case, and each transgene was inserted into the same locus in the *C. elegans* genome (Figure 2, see *Materials and Methods*). We confirmed the knock-ins by observation of the predicted fluorescence localization pattern at the plasma membrane, and we confirmed that knock-ins were single copy by PCR genotyping (Figure 2 and Supplemental Figure 1B).

### Effects of endogenous autofluorescence on fluorescent protein selection

Because single-copy fluorescent transgenes sometimes produce weak fluorescent signal *in vivo,* we quantitatively assessed the endogenous autofluorescence levels of *C. elegans* embryos. We found autofluorescence to be most prominent under 488nm excitation, across a broad range of emission wavelengths (Figure 1A). Thus, when expressed at low levels, fluorescent proteins excited by 488nm light, including GFP, will have poor signal to noise in *C. elegans* embryos. Embryos had considerably less autofluorescent background with 514nm excitation (Figure 1A). This suggests that mNeonGreen may be superior to GFP when imaging single-copy fluorescent proteins that are expressed at low levels in the embryo, given that the microscope setup allows for the excitation of the fluorescent protein with 514nm wavelength.

**Figure 1.**
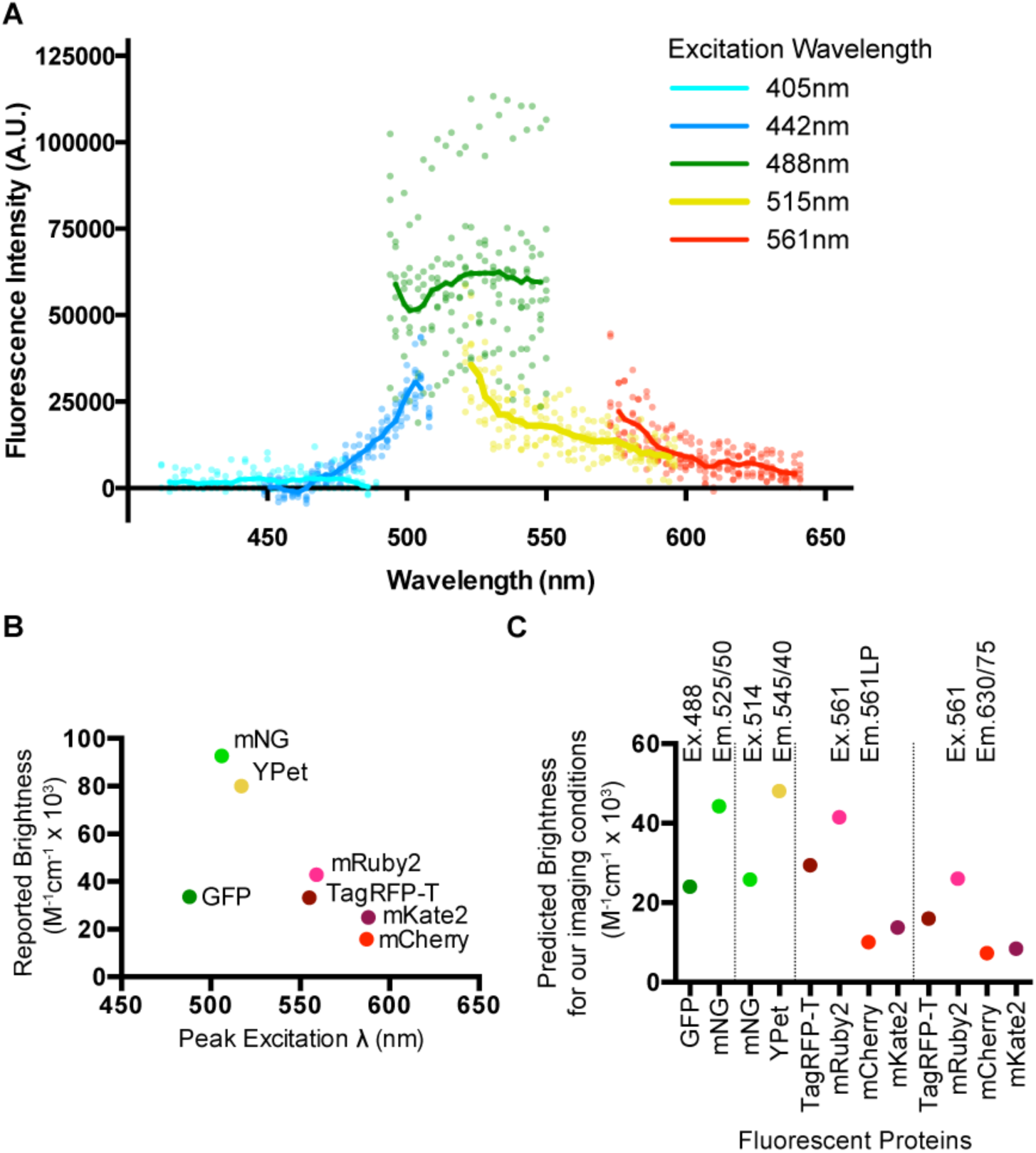
Embryo autofluorescence and predicted brightness of fluorescent proteins. A) Embryo autofluorescence. Lines are averages of multiple embryos. Points are individual embryos. B) Reported brightness for fluorescent proteins at peak excitation wavelengths. C) Predicted brightness of fluorescent protein comparisons performed in Figure 2. Excitation and emission wavelengths are at top.

### *In vivo* fluorescent protein brightness

Before making *in vivo* measurements, we made quantitative predictions about which fluorescent proteins were expected to be brightest. We calculated the potential brightness of each fluorescent protein by taking the product of the quantum yield and extinction coefficient as reported in the literature (Figure 1B)(Yang *et al.,* 1996; Shaner *et al.,* 2004; Nguyen and Daugherty, 2005; Shaner *et al.,* 2008; Shcherbo *et al.,* 2009; Lam *et al.,* 2012; Lee *et al.,* 2013; Shaner *et al.,* 2013). Because imaging conditions such as excitation wavelength and emission filter sets used impact the observed brightness of a fluorescent protein, we sought to use these values to make more useful predictions of fluorescent protein brightness for directly comparing with our results.

To facilitate the visual and quantitative evaluation of fluorescent protein spectra with the specific laser lines and filter sets that are used by us and others, we developed a simple and customizable Microsoft Excel-based tool that we call the Spectrum Viewer (Supplemental File S1). Using this tool, we calculated a predicted brightness for each fluorescent protein by integrating the portion of the fluorescent protein emission peak under our emission filter and multiplying by the quantum yield (Figure 1C). We then used the Spectrum Viewer to plot the normalized absorbance and emission spectra for the fluorescent proteins in our comparisons with the excitation wavelength and emission filter sets we used for imaging (Figure 2 A-D, third column). The Spectrum Viewer is available as a Supplemental File S1.

**Figure 2.**
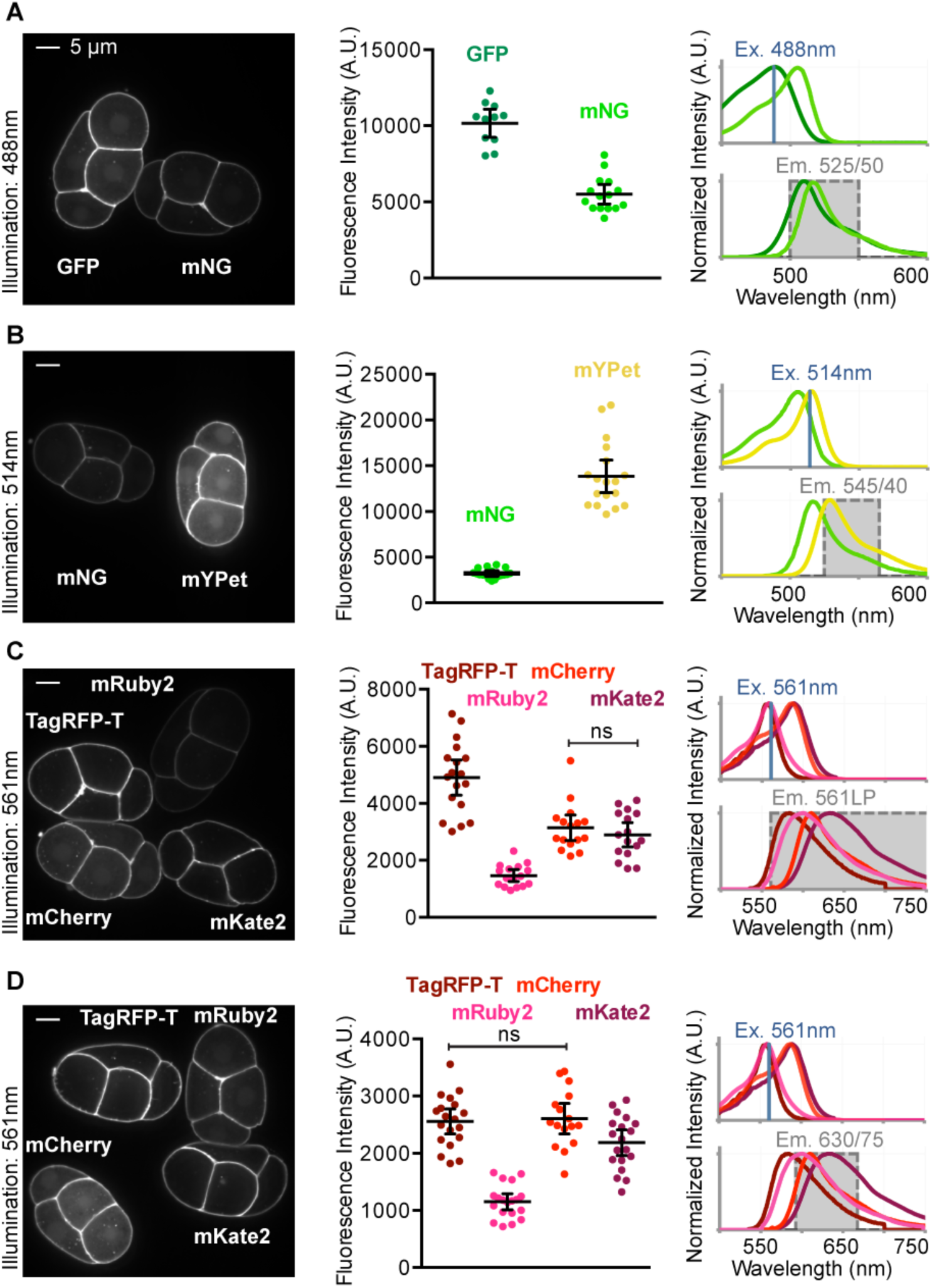
*In vivo* fluorescent protein brightness. (A-D) Left column: Embryos mounted side-by-side and imaged under the same conditions used for quantification. Center column: Graphs show the quantification of each comparison. Each data point represents a single embryo. Black bars indicate the mean and 95% confidence intervals (CIs). Right column: Excitation (upper) and emission spectra (lower) of the compared fluorescent proteins. The illumination wavelength (Ex., blue line) and filter sets used for detection are indicated (Em., gray shading).

To assess the brightness of this set of fluorescent protein transgenes *in vivo,* we imaged staged *C. elegans* embryos, in some cases mounted side-by-side for direct comparisons, by spinning disk confocal microscopy. We first compared GFP and mNG by quantifying the fluorescence from embryos illuminated with 488nm excitation. Although mNG was predicted to be brighter than GFP based on *in vitro* data (Figure 1B and Figure C), we found that the GFP signal was nearly twice as bright *in vivo* (Figure 2A). Mean values within each comparison are significantly different (p<0.05) except where indicated with ns: not significantly different (determined by Student’s t-test with Welch’s correction), and all significance values (P-values) are reported in Supplemental Figure S2, B.

Using 514nm illumination, mYPet was also brighter than mNG (Figure 2B). Though our calculations predicted that mYPet would be almost twice as bright as mNG (Figure 1C), we observed mYPet to be about four times as bright as mNG on average (Figure 2B). The data from the comparisons of mNG with GFP and mYpet suggest that mNG is not as bright *in vivo* as predicted based on the published extinction coefficient and quantum yield (Shaner *et. al.,* 2013) (Figure 1C, Figure 2A and B).

Next we examined the brightness of four red fluorescent proteins (TagRFP-T, mRuby2, mCherry and mKate2). We performed experiments with two different emission filter sets, 561LP and 630/75BP, which are well matched to some or all of these red fluorescent proteins. The 561 LP emission filter is optimal because it collects the majority of the emission peak emission for each fluorescent protein (Figure 2C). A band pass filter, such as the 630/75BP, is less optimal (compare right column Figure 2C and D), however, it may be useful for decreasing spectral overlap for two or three-color imaging.

Using 561 nm illumination we measured the brightness of the four red fluorescent proteins. We found that TagRFP-T was the brightest using the 561 LP filter set (Figure 2C). Using the 630/75BP filter set, the average fluorescence intensity of TagRFP-T was indistinguishable from that of mCherry (Figure 1D). These results are consistent with the orange-shifted emission spectra of TagRFP-T and with our calculated predictions for these fluorescent proteins (Figure 1C, 2C and D). mRuby2, which was predicted to be the brightest of the four red fluorescent proteins (Figure 1C), was the least bright regardless of the emission filter set we used (Figure 2C and D). Taken together, these data reveal fluorescent protein brightnesses *in vivo,* which did not always match predictions made using parameters measured *in vitro*.

### Variation in fluorescent protein brightness between single-copy transgenes

Because we predicted that mNG would be ∼1.8 times brighter than GFP, we were surprised to find that the GFP embryos were significantly brighter than mNG embryos (Figure 1C, 2A). Germline silencing in *C. elegans* can have heterogeneous effects on certain single-copy transgenes (Shirayama *et al.,* 2012). Consequently, fluorescent protein transgenes that are in every other way identical could be expressed at different levels, causing discrepancies between predicted and observed brightness. To ask whether differences in fluorescent protein abundance could account for the differences in fluorescence intensity we observed, we analyzed protein levels in each of our single-copy transgenic strains by western blot (Supplemental Figure S1). We observed approximately 2fold higher levels of *mex-5* driven GFP::PH protein compared with mNG::PH protein (Supplemental Figure S1, C, paired t-test, p=0.0408), which may be due to partial transgene silencing or post-transcriptional regulation of these transgenes.

To further investigate the discrepancy between our predictions and observations, we compared a second set of identical GFP and mNG single-copy transgene knock-in strains. These fluorescent proteins were fused to the C-terminus of a histone gene *(his-58)*. As expected, the resulting fluorescence was brightest in nuclei (Figure 3A). To control for effects of cell cycle timing on histone protein abundance, we staged embryos to within 3 minutes of one another. We measured the fluorescence intensity in the nucleus of one embryonic cell (the EMS cell) in each embryo and found that the average fluorescence intensity of the GFP-histone expressing embryos and the mNG-histone embryos were not significantly different (Figure 3A). Although in our initial comparison of membrane localized transgenes we found that GFP expressing embryos were significantly brighter than those expressing mNG (Figure 2A), both results suggest that in early *C. elegans* embryos mNG is not as bright when compared with GFP as we had predicted (Figure 1C).

**Figure 3.**
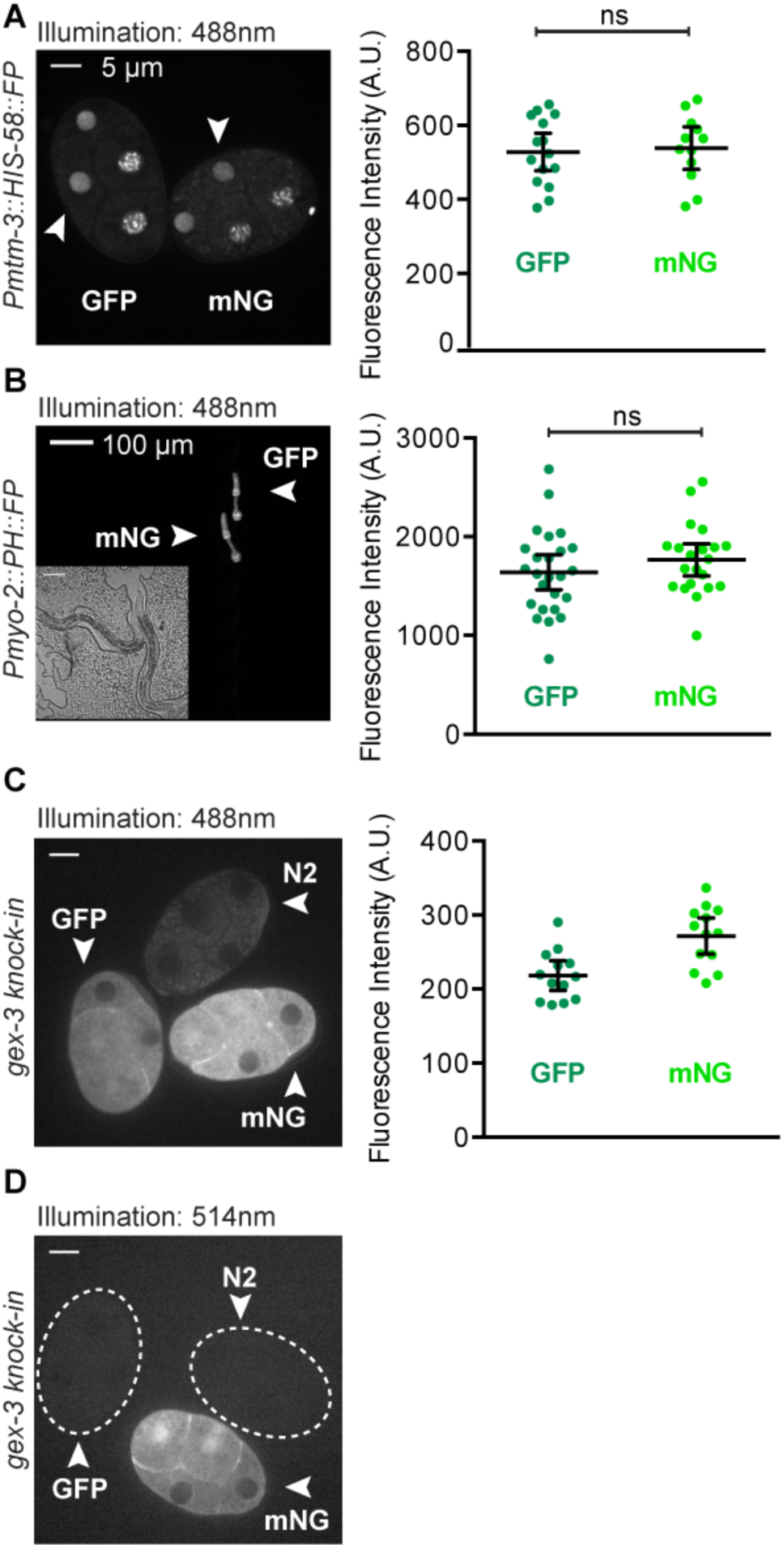
Comparing GFP and mNeonGreen across different tissues. (A-C) Each data point represents a single embryo or animal, black bars represent the mean and 95% CIs. (A) Embryos expressing histone-fluorescent protein fusions. Fluorescence intensity of the EmS cell nucleus was measured (white arrowheads). (B) Young adult worms expressing membrane tag-fluorescent protein fusions in the pharynx (white arrowheads). The insert is a DIC image of the worms. (C and D) GFP::GEX-3, mNG::GEX-3, and N2 wild type embryos were imaged using (C) 488nm illumination and (D) 514nm illumination. Dotted lines outline GFP::GEX-3 and N2 wild type embryos.

Because protein levels in the *C. elegans* germline and early embryo can be affected by silencing mechanisms (Shirayama *et al.,* 2012), we compared GFP and mNG in a *C. elegans* tissue that has not been reported to exhibit the silencing. We replaced the germline promoter in our original GFP and mNG::PH repair template constructs with the *myo-2* promoter, which drives expression in the pharynx (Okkema *et al.,* 1993) and generated single-copy transgene knock-ins at the same genomic locus used for our initial comparison. We imaged staged worms and quantified GFP and mNG fluorescence and again found no significant difference between average GFP and mNG intensities (Figure 3B). These data are consistent with our findings in early embryos, and are consistent with the possibility that factors outside of germline silencing play a role in determining the observed fluorescence from single-copy transgenes.

### Comparing green fluorescent proteins as endogenous tags

We next set out to compare GFP and mNG at lower levels of expression and inserted into an existing gene at its endogenous locus. Previous work had shown that an N-terminal mNG knock-in at the *gex-3* locus (a member of the actin regulatory WAVE complex) was detectable in the early embryo but not highly expressed (Dickinson *et. al.,* 2015). We created an identical GFP::GEX-3 strain for comparison. We imaged embryos from the two strains side-by-side at the same developmental stage as in our previous comparisons. Using 488nm illumination, on average, mNG::GEX-3 embryos were brighter than GFP::GEX-3 embryos (Figure 3C).

Because background embryo autofluorescence is higher at 488nm illumination (Figure 1A), we also imaged these embryos using 514nm illumination. Although we could not quantitatively compare fluorescence intensity of embryos illuminated with 488nm vs. 514nm wavelengths due to differences in image acquisition set-up (e.g. laser power, filter sets, etc.), we observed that mNG imaged with 514 nm illumination gave the clearest picture of the actual fluorescent protein signal due to the lower background autofluorescence (Compare mNG vs. N2 signal in Figures 3C and D).

### Photostability of fluorescent proteins *in vivo*

The brightness of a fluorescent protein together with its photobleaching rate determine how useful a fluorescent protein is for time-lapse imaging (Shaner *et. al.,* 2005; Davidson and Campbell, 2009; Shaner, 2014). To test the rate of photobleaching of the fluorescent proteins used in our initial comparison in Figure 2, we imaged embryos over time under continuous illumination (Figure 4 A-C). Fluorescence intensities were normalized to initial brightness measured for each embryo, and averages were plotted for each strain over time (Figure 4 A-C; left). Each photobleaching curve was fit to a one-phase exponential decay and the half-life was calculated (Figure 4 A-C; middle). To estimate a “photon-budget”, or the amount of signal emitted by each fluorescent protein over time, we integrated the fluorescence intensity measured for each embryo up to 50% of its initial intensity (Figure 4 A-C; right) (Lee *et. al.,* 2013).

**Figure 4.**
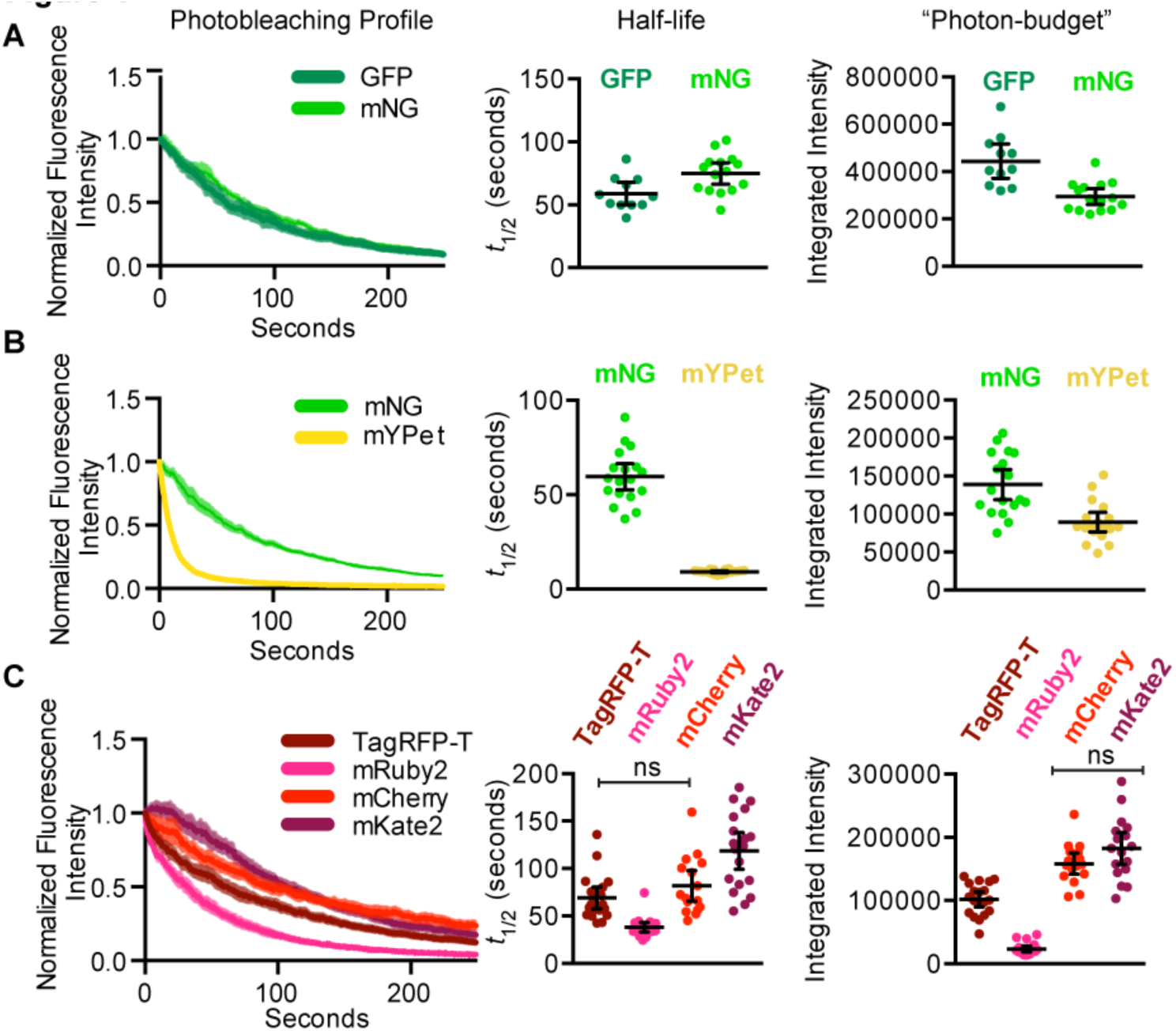
*In vivo* fluorescent protein photostability. (A-C) Fluorescence intensity was measured in embryos over time. Photobleaching profile, half-life and photon-budget were compared for membrane-fluorescent protein fusions. Each data point represents a single embryo, and the black bars represent the mean and 95% CIs.

GFP and mNG displayed similar bleaching half-life, with mNG being slightly more photostable (Figure 4A). However, because the GFP embryos are brighter, on average, the integrated intensity, or photon-budget of the gFp embryos was slightly higher than that of mNG (Figure 4A). mYPet was observed to bleach far faster than mNG, as expected (Figure 3B) (Shaner *et. al.,* 2013). Of the red fluorescent proteins we tested, mKate2 had the slowest average bleaching rate (Figure 3C). The photobleaching profile of mKate2 suggests that it exhibits kindling (photoactivation) phenomenon in the first few frames of illumination (Figure 3C and Supplemental Figure S3). Photoactivation was not reported in the initial characterization of mKate2, but had been observed for its precursor protein mKate (Shcherbo *et al.,* 2009). We conclude that mRuby2 and mYPet exhibited relatively poor photostability *in vivo,* and that GFP, mNG and mKate2 were most photostable.

### Summary and Recommendations

Our results suggest specific recommendations for fluorescent proteins to use in *in vivo* experiments in *C. elegans* embryos, forming a baseline for comparisons in other *in vivo* systems. In general, we observed a lower-than-expected brightness for mNG. Using 488nm illumination, GFP and mNG performed similarly: in two comparisons GFP and mNG were equally bright, GFP was brighter in a germline transgene expressed at high levels, and mNG was brighter as a knock-in at an endogenous gene locus (Figure 2A and Figure 3). mYPet was significantly brighter than mNG, but its high rate of photobleaching makes it an unattractive choice for imaging more than a few frames (Figure 2B and Figure 4B). Of the red fluorescent proteins we tested, we recommend using mKate2. Although TagRFP-T, mCherry, and mKate2 performed similarly in terms of brightness, mKate2 had the superior photobleaching dynamics *in vivo* (Figure 2C, D and 4C).

Our measurements of autofluorescence in the early *C. elegans* embryo highlight the value of taking such measurements before designing *in vivo* imaging experiments. For *C. elegans* embryos, using 488nm illumination will give higher background than imaging using 514nm illumination (Figure 1A). Therefore, for genes with low expression levels, better signal-to-noise ratios may be achieved using a yellow fluorescent protein and exciting with 514nm illumination, rather than a green fluorescent protein and 488nm illumination (Figure 3C and D). Because of the rapid photobleaching we observed for mYPet (Figure 4B), we would choose mNG to tag proteins expressed at low levels in *C. elegans* embryos for live-cell imaging.

It is unknown how applicable the specific results of this study are in model systems beyond *C. elegans*. The fluorescent proteins we found to be optimal were not the same as another comprehensive *in vivo* comparison of fluorescent proteins in yeast (Lee *et. al,* 2013). Variables such as autofluorescence and gene silencing may influence fluorescent protein choice to a greater or lesser extent across diverse cell types and model systems. Future studies in diverse systems are needed to reveal whether there is a universally best set of fluorescent proteins. We used exclusively spinning disk confocal microscopy for our comparison. However, differences in illumination source and detectors used in different light microscopy techniques (e.g. widefield, TIRF, lightsheet, etc.) may change the observed performance of fluorescent proteins in live-imaging experiments.

This study contributes information of practical value about which fluorescent proteins to use for *in vivo* experiments, as well as a tool for researchers to use to evaluate the spectra of different fluorescent proteins relative to their own imaging resources. The findings are especially applicable for experiments in *C. elegans,* and they suggest the value of performing similar experiments in other model systems.

## Materials and Methods

### *C. elegans* Strains and Maintenance

All *C. elegans* strains used in this study are listed in Supplemental Figure S1 and were handled using standard techniques (Brenner, 1974). The strains were raised at 25°C, in incubators in the dark, and fed *E. coli* 0P50 except where otherwise indicated. The HT1593 *(unc-119(ed3)* III) strain, used as the parent to the LP306, LP274, LP402, LP193, LP307, LP308, LP401, LP403, and LP404 strains generated in this study, was raised at 15°C and fed *E. coli* HB101 prior to injection (Hochbaum *et al.,* 2010; Dickinson *et al.,* 2013).

### Fluorescent Protein Selection

Because of their current widespread use, we chose to compare GFP and mCherry with newer green and red fluorescent proteins that are less commonly used but that have been described as having superior brightness and/or photostability. We used a GFP variant, GFP S65C, commonly used in *C. elegans,* which we will refer to as GFP (Green *et al.,* 2008). S65C and S65T (eGFP) variants perform similarly (Heim and Tsien, 1996), and a previous *in vivo* study of fluorescent proteins in yeast reported that S65T outperformed certain green fluorescent protein variants (such as Clover and Emerald) in a direct comparison (Lee *et al.,* 2013). mNeonGreen (mNG), is a newer, monomeric green fluorescent protein (peak excitation ∼506nm) that is reported to be up to three times as bright and more photostable than eGFP *in vitro* (Shaner *et al.,* 2013). We therefore compared mNG to GFP in our *in vivo* system. To assess the practical value of mNG’s yellow-shifted excitation spectrum (Shaner *et al.,* 2013), we compared mNG with a yellow fluorescent protein, mYPet—the brightest reported yellow fluorescent protein reported to date (Nguyen and Daugherty, 2005). We chose three red fluorescent proteins to compare with mCherry: TagRFP-T, mKate2, and mRuby2. A direct comparison in yeast found that all three were brighter than mCherry *in vivo* (Lee *et al.,* 2013). These red fluorescent proteins range in peak emission from 584nm to 633nm (Shaner *et al.,* 2008; Shcherbo *et al.,* 2009; Lam *et al.,* 2012), making them useful in combination with different fluorescent proteins for two‐ or three-color imaging.

### Fluorescent protein optimization and repair template construction

Single-copy transgenic knock-in strains (LP306, LP274, LP402, LP193, LP307, LP308, LP401, LP403, LP404) were generated using the method described in Dickinson *et. al.,* 2013. Fluorescent protein sequences were obtained from the following sources ((Heim and Tsien, 1996; Shaner *et al.,* 2004; Nguyen and Daugherty, 2005; Shaner *et al.,* 2008; Shcherbo *et al.,* 2009; Lam *et al.,* 2012; Shaner *et al.,* 2013). We used a mutant form of GFP, S65C, commonly used in *C. elegans*. mNeonGreen was licensed from Allele Biotechnology. To increase the monomeric character of YPet, we introduced a well-characterized mutation to the original YPet sequence (A206K) to generate mYPet (Zacharias *et al.,* 2002; Ohashi *et al.,* 2007).

Repair template constructs were identical, except for the sequences of the fluorescent proteins tested. Each transgene construct consisted of a germline promoter sequence *(Pmex-5)* driving the expression of a fluorescent protein fused to the N-terminus of the same polypeptide: the pleckstrin homology domain from phospholipase C-δ1 (PH domain) and a 2x Flag epitope tag. The PH domain localizes to the plasma membrane by binding phosphatidylinositol 4,5-bisphosphate (PIP^2^) (Audhya *et. al.,* 2005). Because codon usage bias affects the expression level of genes in *C. elegans,* the nucleotide sequences of the fluorescent proteins and PH domain were optimized for expression in *C. elegans* using the *C. elegans* Codon Adapter (CAI ∼1) (Redemann *et al.,* 2011). Synthetic *C. elegans* introns were added to each fluorescent protein to facilitate expression of the transgenes (Fire *et.al.,* 1990). The fluorescent protein genes were synthesized in ∼500bp overlapping gBlock fragments (Integrated DNA Technologies), assembled using Gibson Assembly Master Mix (NEB), PCR amplified, and cloned using the Zero Blunt TOPO PCR cloning kit (Invitrogen).

All repair template constructs were made using a derivative of the pCFJ150 vector backbone modified for Cas9 mediated homologous recombination (Frøkjær-Jensen *et al.,* 2008; Dickinson *et al.,* 2013). The *mex-5* promoter, the *C. elegans* sequence-optimized mNeonGreen fluorescent protein and PH domain, and the *tbb-2* 3’UTR were added using Gibson Assembly (NEB) to create vector pAP006. To generate repair templates with different fluorescent protein sequences, pAP006 was amplified into a linear fragment using the forward primer 5’ CACGGACTCCAAGACGAC (binds after the *mex-5* promoter) and reverse primer 5’ TCTCTGTCTGAAACATTCAATTGATTATC (binds at the start of the *C. elegans* optimized PH domain). Fluorescent protein genes were amplified using gene-specific primers with minimum 30bp overlapping sequence to the parent vector fragment (Forward 5’ C GAT AAT CAATT GAAT GTTT CAGACAGAGA + FP sequence; Reverse 5’ GCCGGCCACGGACTCCAAGACGACCCAGACCTCCAAG + FP sequence). The vector backbone fragment and fluorescent protein genes were assembled using Gibson Assembly (NEB). The repair templates for strains LP403 and LP404 were made using a similar strategy to exchange the *mex-5* promoter for the *myo-2* promoter sequence.

We have deposited constructs containing the optimized fluorescent proteins in Addgene.

### Insertion and confirmation of transgene knock-ins

Single-copy transgenes were inserted into the *C. elegans* genome via Cas9 triggered homologous recombination, using the reagents and methods described in Dickinson *et. al.,* 2013. The transgenes were inserted near the *ttTi5605* MosI insertion site on *C. elegans* chromosome II. This site has been used for both CRISPR/Cas-9 and *Mos1* transposon-based transgene insertions and is known to permit the expression of transgenes in the germline (Frøkjær-Jensen *et al.,* 2008; Dickinson *et al.,* 2013). We used a guide RNA with the following target sequence: 5’ - GATATCAGTCTGTTTCGTAA (Dickinson *et al.,* 2013). Singlecopy knock-ins were confirmed by rescue of the HT1593 uncoordinated phenotype, observation of the predicted fluorescence localization pattern at the plasma membrane, and PCR genotyping (Figure 2 and Supplemental (Figure 1, B). PCR genotyping was performed on genomic DNA extracted from putative knock-in animals, using primers outside the insertion site (5’ - AGGCAGAATGTGAACAAGACTCG and 5’ - ATCGGGAGGCGAACCTAACTG) as described in Dickinson *et. al.,* 2013. All seven transgenes resulted in minimal embryonic lethality at 25°C (Supplemental Figure S2, A).

JA1699 was made using standard MosSCI methods using pJA449 (*mtm-3* associated HOT core/fr/’s-58/mNeonGreen::tbb-2 3’UTR), which was constructed using triple gateway into pCFJ150 using *mtm-3* promoter in pDONRP4P1R, pJA273 *(his-58* coding in pDONR221) and pJA448 *(C. elegans* optimized mNeonGreen::tbb-2 3’UTR in pDONR P2R-P3) (Zeiser *et al.,* 2011; Dickinson *et al.,* 2013). The construction of strain JA1610 is described in Chen *et. al.,* 2014 (Chen *et al.,* 2014). LP431 (GFP::gex-3) was made using the strategy described for LP362 (mNG::gex-3) in Dickinson *et. al.,* 2015 (Dickinson *et. al.,* 2015). PCR genotyping of LP431 was performed using the following primers: 5’ - aactgccgccaacaaaagag and 5’ - ctcacCGCCGCTTGATT.

### Predicted Brightness Calculation

The predicted brightness for each fluorescent protein on our imaging set-up was calculated by taking the sum of the normalized emission spectra over the range of the filter set used for imaging (calculated from our Spectrum Viewer) and dividing by the sum of the normalized emission spectra over the entire spectrum (Spectrum Viewer), and then multiplying by the brightness. The brightness was calculated as a product of the quantum yield and the extinction coefficient times the fraction of excitation peak at the imaging wavelength (Yang *et al.,* 1996; Shaner *et al.,* 2004; Nguyen and Daugherty, 2005; Shaner *et al.,* 2008; Shcherbo *et al.,* 2009; Lam *et al.,* 2012; Lee *et al.,* 2013; Shaner *et al.,* 2013).

## Microscopy

### Imaging embryos

*C. elegans* embryos were dissected for imaging and mounted in egg buffer at 2-3 cell stage on poly-L-lysine coated coverslips, with 2.5% agar pads. Embryos expressing different fluorescent proteins were initially imaged side-by-side, as shown in Figures 2 and 3 (n=3 pairs/groups per comparison). To increase the number of embryos imaged for quantification, multiple embryos from the same strain were mounted in groups and images were acquired using the same settings as the initial side-by-side comparisons. To minimize the effect of any unavoidable, minor, variation in imaging conditions, embryos from strains for a given comparison were imaged alternately using identical settings. HIS-58::GFP and mNG embryos were mounted at the three-cell stage, a short (∼3min), identifiable stage between cell divisions. Fluorescence intensity was measured in the EMS cell nucleus. For the GFP and mNeonGreen::GEX-3 knock-in strain comparisons, embryos from each strain plus an N2 wildtype embryo were imaged and compared in groups.

All embryos were imaged with a Nikon Eclipse Ti spinning disk confocal microscope (Yokogawa CSU-X1 spinning disk head) using a Hamamatsu ImagEM X2 EM-CCD camera (C9100-13) and a 60X/1.4 NA Plan Apo oil immersion objective (Nikon). Samples were illuminated using solid-state lasers of the following wavelengths: 488nm, 514nm, and 561nm. The following emission filter sets were used for a given excitation wavelength: 488nm: ET525/50m (Chroma), 514nm: ET545/40m (Chroma), 561nm: ET630/75m (Chroma) and 561 lp (Semrock).

### Imaging whole worms

Whole worms were mounted at the L4 developmental stage and immobilized using nano-particles as previously described (Kim *et al.,* 2013). Worms were imaged using a Nikon Eclipse Ti spinning disk confocal microscope (Yokogawa CSU-X1 spinning disk head) using a Hamamatsu ImagEM X2 EM-CCD camera (C9100-13) and a 10X/0.30 NA Plan Fluor objective (Nikon) with 488nm excitation and ET525/50x emission filter.

### Image Quantification

For membrane labeled strains, fluorescence intensity was quantified using Metamorph Software (Molecular Devices) by taking the average of a 3 pixel wide linescan perpendicular to the plasma membrane in the posterior-most embryonic cell (the P^2^ cell). For each time point, the maximum intensity from this linescan was recorded and average off-embryo background was subtracted. GraphPad Prism software was used to plot the mean and 95% confidence intervals (CIs) for all initial brightness measurements, and at each time point for bleaching measurements. To determine the half-life of a given fluorescent protein, the individual photobleaching traces were fit to a standard one-phase decay curve, the ‘half-life’ for each curve was recorded, and the mean and 95% CIs were recorded for each fluorescent protein. The photon-budget was determined by integrating the fluorescence intensity measured for each embryo until the intensity reached 50% of the initial intensity.

For histone fusion proteins and pharyngeal labeled strains, the images were thresholded and segmented using ImageJ to define a region for measurement (either the nucleus or pharynx). For GFP and mNeonGreen::GEX-3 knock-in strains, a region was drawn around each embryo. The average fluorescence intensity of the given regions were calculated by measuring the average integrated intensity of the region and subtracting average off-embryo background for each image. Each embryo was displayed as an individual data point, and the mean and 95% CIs were plotted using GraphPad Prism software.

Unpaired, two-tailed t-tests with Welch’s correction were used to compare means in all imaging experiments, and all statistical analyses were performed using GraphPad Prism software (GraphPad Software, Inc.). All comparisons are significantly different (p<0.05), unless otherwise indicated by ‘ns’. Statistics for individual experiments can be found in Supplemental Figure S2, B.

### Quantifying auto-fluorescence in *C. elegans* embryos

N2 (wild type) embryos were mounted in egg buffer on poly-l-lysine coated coverslips, with 2.5% agar pads. The embryos were imaged using a Nikon A1R laser scanning confocal microscope. The excitation wavelengths used were 405nm, 442nm, 488nm, 515nm, and 561nm. The illumination settings for each wavelength were set to a common wattage in the Nikon elements software. Images of embryo autofluorescence were collected using a multispectral detector and emission fingerprinting for each of the given wavelengths. Image analysis was performed using ImageJ. Pixel intensity values were measured for three regions per embryo and averaged. Average off-embryo background was subtracted for each embryo, and the resulting fluorescence intensity was plotted at each detection wavelength.

### Western blotting

For quantifying protein levels, L4 staged worms were picked to plates 12-14 hours prior to lysis. Lysates were generated at a concentration of one worm per microliter (60 worms were picked into 45μl M9 Buffer and15μl 4X Sample Buffer was added). Samples were frozen in liquid nitrogen and sonicated in boiling water for 10 minutes twice. Lysates were separated on 12% NuPAGE Novex Bis-Tris Protein Gels (Invitrogen) and transferred to an Immobilon PVDF-FL membrane (Millipore) for immunoblotting. Fluorescent proteins expressed by transgenes were detected using a mouse anti-FLAG BioM2 (Sigma-Aldrich, catalog number F9291) antibody at 1:1000 dilution, and a rabbit anti-HCP-3 (Monen *et. al.,* 2005) was used at 1:1000 dilution as a loading control. The following fluorescent secondary antibodies were used (1μl per blot): AlexaFluor 680 goat anti-mouse and AlexaFluor 790 goat anti-rabbit (Invitrogen catalog numbers, A31562 and A11369, respectively). Samples were collected and blots were performed in triplicate. Blots were scanned using an Odyssey Infrared Imaging System (LI-COR Biosciences) and fluorescence intensity was quantified using ImageJ. A ratio of transgene protein intensity (∼45kDa band in 680nm channel) to loading control intensity (upper band in 790nm channel) was measured for each lane on a given blot. These measurements were normalized by dividing by the ratio measured for each lane, by the total average ratio of all the lanes on a given blot. These normalized protein levels were plotted along with an average and 95%CIs using GraphPad Prism. Gel images were inverted and cropped slightly at the edges, and brightness and contrast were adjusted using ImageJ. Dashed line indicates were blank lanes were cropped.

### Spectrum Viewer

The fluorescence spectrum viewer (Supplemental File S1) was designed as a user-extensible collection of fluorescence spectra, dichroic filter spectra, and laser lines. Data was collected and digitized from a range of published fluorophore spectra using the WebPlotDigitizer software package (http://arohatgi.info/WebPlotDigitizer). Digitized spectra were resampled at one nanometer wavelength increments and excitation and emission spectra were each normalized to a maximum value of one relative fluorescent unit. Dichroic fluorescence filter data were similarly digitized from commercial plots. The spectrum viewer was implemented in Microsoft Excel using only worksheet range functions and avoids the use of macro-language constructs. Up to four fluorophores, four fluorescent filters, and three laser lines may be selected and compared in an Excel chart through a simple graphical user interface. Possible spectral data listed in the user interface are populated from a “DataList” database worksheet, which in turn consists of spectrum names and accompanying worksheet ranges for stored spectral data. User selection of a spectrum to display populates a “Current” data worksheet via indirect references stored in the “DataList” database. The spectral chart is automatically updated to reflect changes in the “Current” data worksheet.

New fluorophore and fluorescent protein spectral data may be added to existing worksheets or as new worksheets. Indirect worksheet references must then be added to either the fluorophore or filter section of the “DataList” worksheet. The user interface is automatically repopulated with new choices. Simple, user-defined bandpass, shortpass, and longpass filter sets may also be defined on the “User Filters” worksheet for comparison to fluorophore spectra.

## Acknowledgements

We thank Kurt Thorn, Dave Matus, and Michael Werner for suggestions and comments on the manuscript. We also thank Terrence Wong for materials and plates. Strains were provided by the Caenorhabditis Genetics Center, which is funded by the U.S. National Institutes of Health (NIH) Office of Research Infrastructure Programs (P40 OD010440). This work was supported by NIH T32 CA009156 (D.J.D. and A.M.P.); a Howard Hughes postdoctoral fellowship from the Helen Hay Whitney Foundation (D.J.D.); a U.S. National Science Foundation (NSF) graduate research fellowship (J.K.H.); and NIH R01 GM083071 and NSF IOS 0917726 (B.G.).

## Author Contributions

Conceived and designed experiments: J.K.H., D.J.D., A.M.P., C.D.H., B.G.; Performed experiments: J.K.H.; Analyzed Data: J.K.H.; Drafted the Article: J.K.H., B.G.; Prepared the digital images: J.K.H.; Constructed and contributed reagents: J.K.H., D.J.D., A.M.P., C.D.H., A.S., J.A.; Developed the spectrum viewer: J.R.K.

---

**Supplemantal Figure 1.**
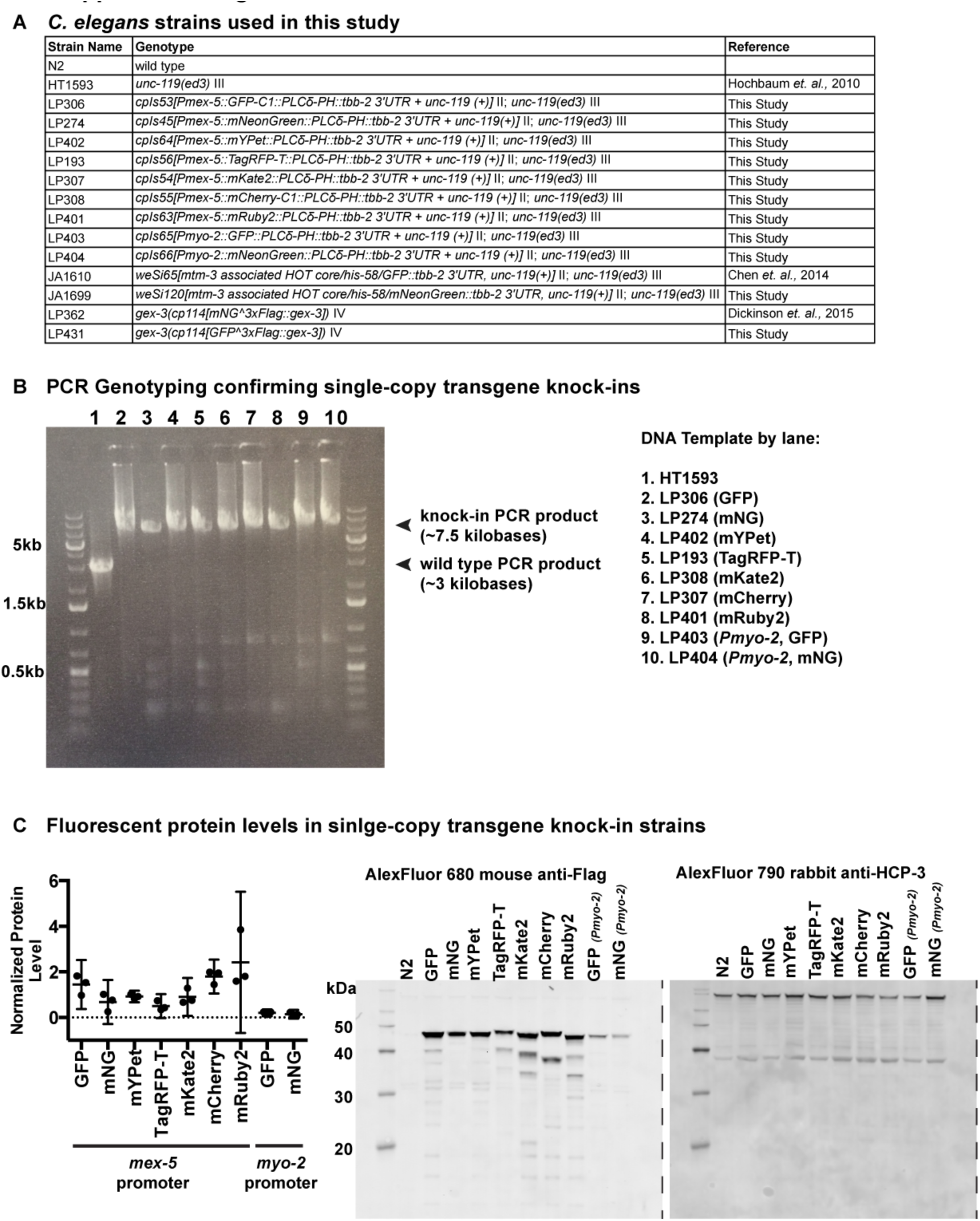
A) *C. elegans strains* A list of all the *C. elegans* strains used in this study. B)PCR genotyping confirming single-copy transgene knock-ins PCR genotyping was performed using primers that flank the Cas9 target site on *C. elegans* chromosome II. The increased size (+4.5kb) of the PCR products in lanes 2-10 indicate a single-copy insertion. C) Fluorescent protein levels in single-copy transgene knock-ins Lysates from worms expressing FP::PH::2XFlag driven by either the *mex-5* (embryos) or *myo-2* (pharynx) promoter were immunoblotted. Individual data points are normalized protein levels for each strain and black bars are a mean and 95% CIs.

**Supplemantal Figure 2.**
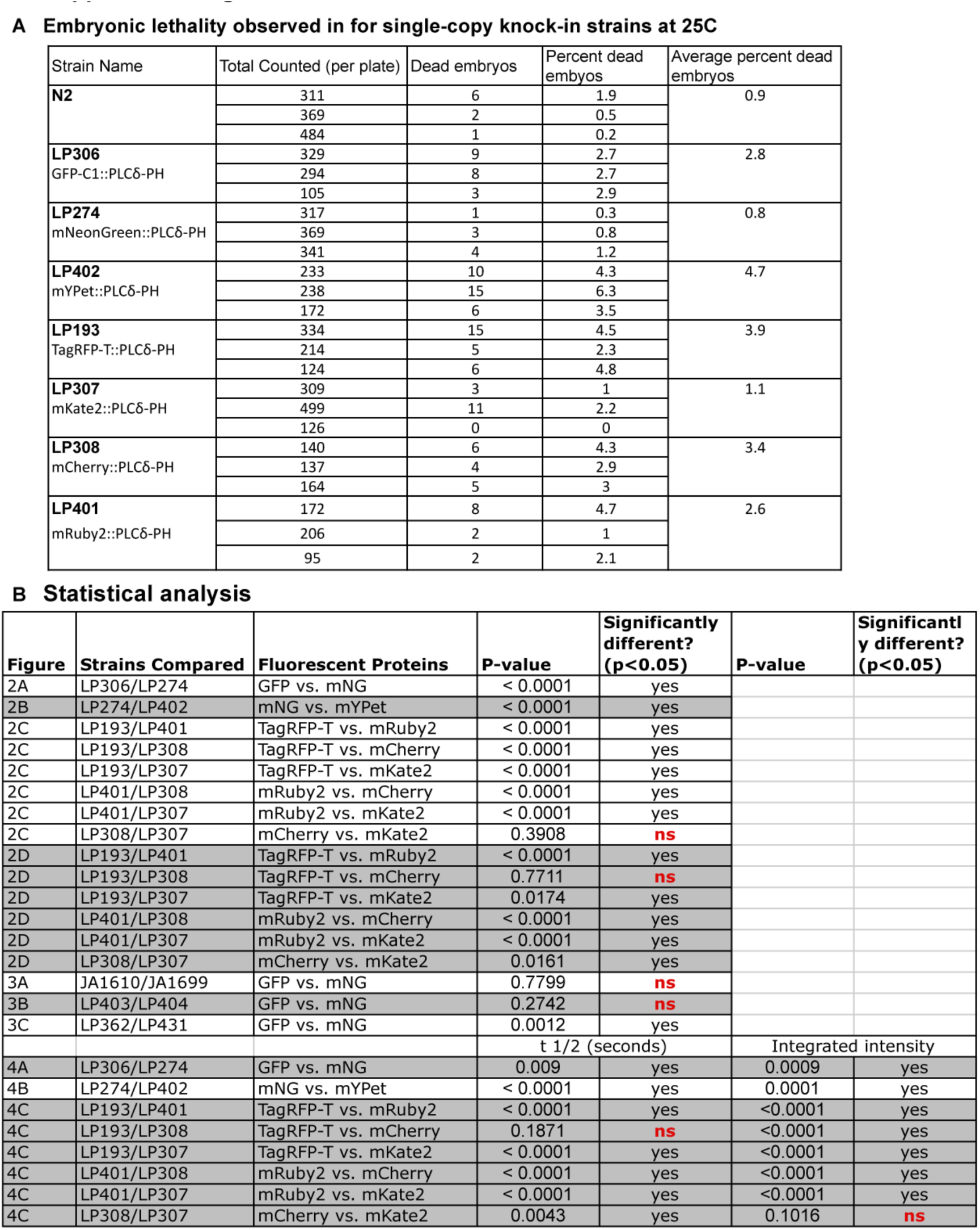
A) Embryonic lethality B) Statistical analysis The calculated P-value was judged as significantly different (p<0.05, yes) or not significantly different (p>0.05, ns). Non-significant results are labeled in the main text figures.

**Supplemantal Figure 3.**
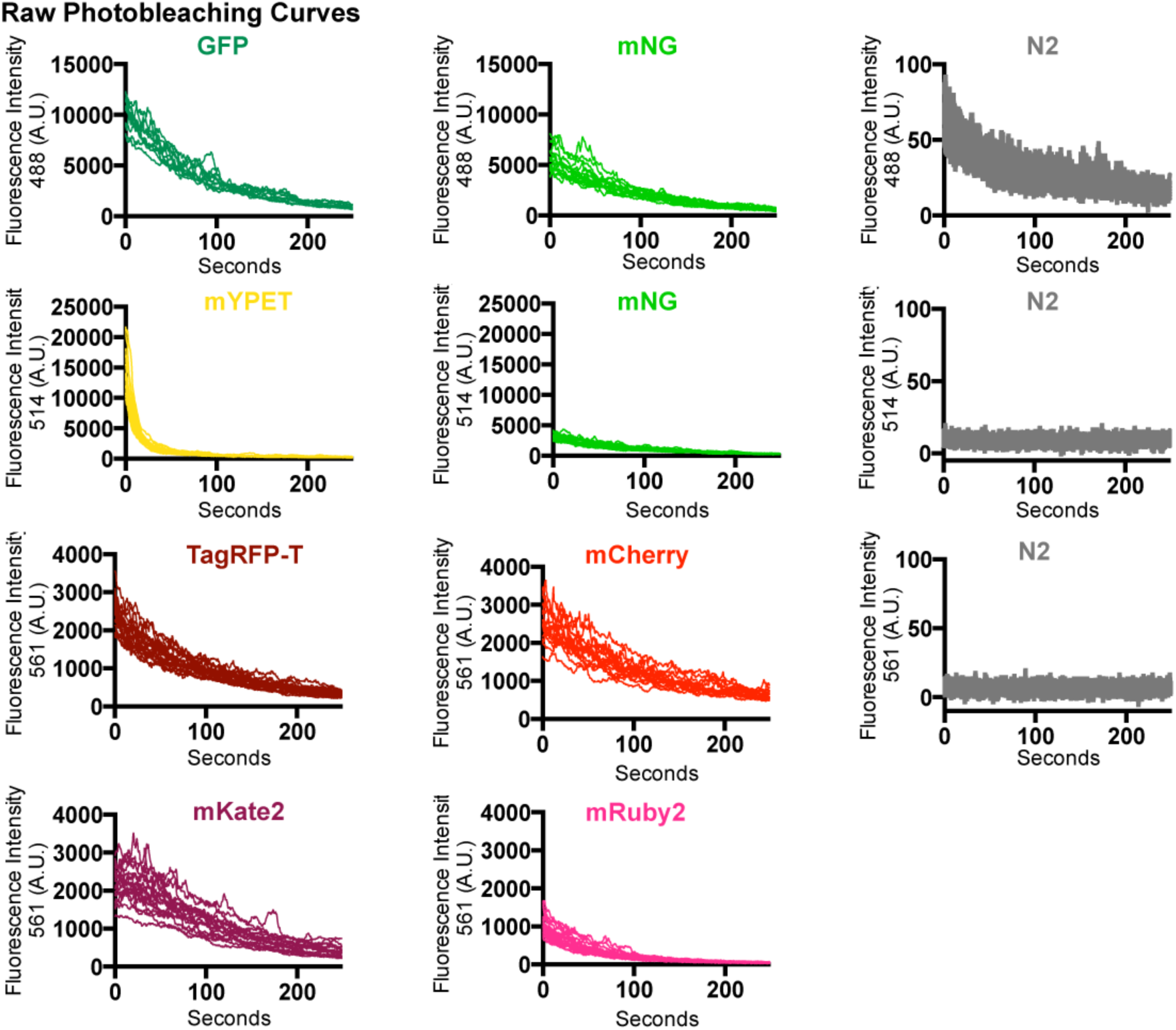
Raw photobleaching curves

